# Spatially coordinated collective phosphorylation filters spatiotemporal noises for precise circadian timekeeping

**DOI:** 10.1101/2022.10.27.513792

**Authors:** Seok Joo Chae, Dae Wook Kim, Seunggyu Lee, Jae Kyoung Kim

## Abstract

The circadian (∼24h) clock is based on a negative feedback loop centered around the PERIOD protein (PER), translated in the cytoplasm and then enters the nucleus to repress its own transcription at the right time of day. Such precise nucleus entry is mysterious because thousands of PER molecules transit through crowded cytoplasm and arrive at the perinucleus across several hours. To understand this, we developed a mathematical model describing the complex spatiotemporal dynamics of PER as a single random time delay. We find that the spatially coordinated bistable phosphoswitch of PER, which triggers the phosphorylation of accumulated PER at the perinucleus, leads to the synchronous and precise nuclear entry of PER. This leads to robust circadian rhythms even when PER arrival times are heterogenous and perturbed due to changes in cell crowdedness, cell size, and transcriptional activator levels. This shows how the circadian clock compensates for spatiotemporal noise.

**Graphical Abstract:** 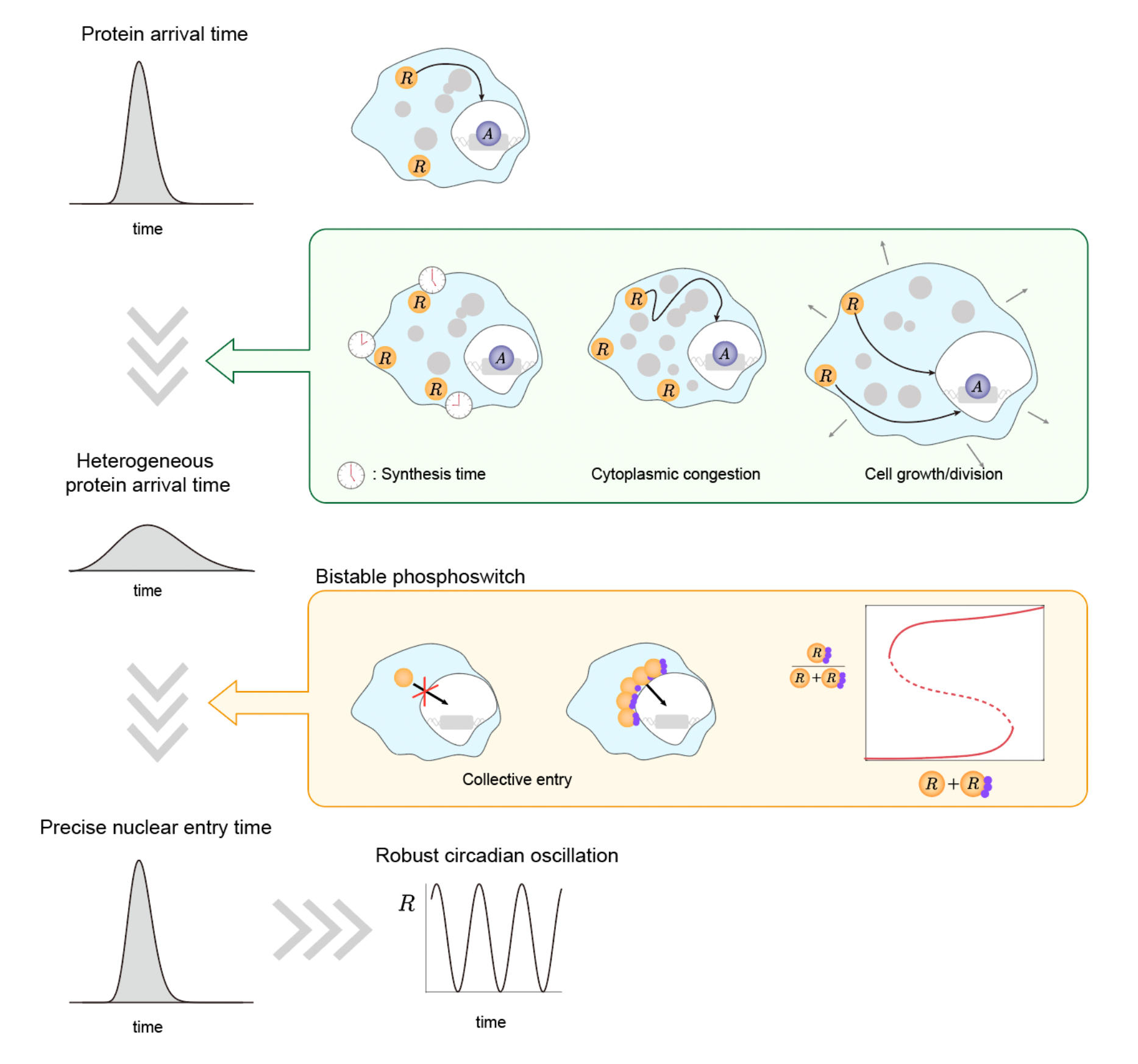

**Highlights:**

- The time window when PER protein arrives at the perinucleus is wide and keeps changing.
- A bistable phosphoswitch enables precise nuclear entry of PER protein.
- This leads to robust circadian rhythms when cell congestion level and size change.
- This describes how the circadian clock compensates for spatiotemporal noise.

## Introduction

The mammalian circadian (∼24h) clock is a self-sustained endogenous oscillator that relies on a transcriptional-translational negative feedback loop (TTFL), where the activator complex, BMAL1:CLOCK, promotes the transcription of *Per1/2* and *Cry1/2* genes, and the PER:CRY complex inhibits BMAL1:CLOCK to close the loop^1-6^. In the TTFL, precise transcriptional repression (i.e., transcription being repressed at the right time of the day) of BMAL1:CLOCK by the PER complex is essential to generate circadian rhythms^7-10^.

However, obtaining such precise transcriptional repression is challenging because individual PER molecules are expected to arrive at the perinucleus at different times. Specifically, thousands of PER molecules transit through the crowded intracellular environment with organelles and macromolecules, leading to different travel times for each PER molecule. Moreover, since the cell size keeps changing due to cell growth and cell division, the travel distance of PER molecules to the perinucleus also keeps changing. The heterogeneity in the PER arrival time is further amplified as PER molecules are translated across several hours at different places^8,11,12^. Furthermore, the amount of activator proteins of the transcription (e.g., BMAL1), which promotes the production of PER protein, also exhibits a noisy fluctuation with daily changes^13,14^. As a result, the arrival time of PER molecules largely varies during a single day and exhibits daily variations. Interestingly, although thousands of PER molecules arrive at the perinucleus across several hours, they enter the nucleus during a narrow time window at the right time every day^8^, leading to transcriptional repression with precise timing, and thus to robust circadian rhythms. This indicates the existence of some mechanism that filters the heterogeneity in the protein arrival time.

This filtering mechanism has barely been investigated. Even the widely used mathematical models of the circadian clock assume that PER is homogenously distributed in the cytoplasm, and thus they are not able to capture the cytoplasmic trafficking of PER^15-31^. Notably, it has been shown that when the cytoplasmic trafficking of Hes1 molecules is incorporated into a mathematical model, the period of Hes1 oscillation greatly changes (∼three-fold)^32^. This indicates that the variability in the protein arrival time distribution should be filtered to generate stable rhythms. Previously, a potential filtering mechanism was proposed: If traveling molecules degrade quickly enough, molecules that spend a long time during cytoplasmic trafficking are degraded before they arrive at the perinucleus^33^, and thus only molecules traveling along optimal paths enter the nucleus without being degraded. Although this results in a narrow distribution of nuclear entry time, it requires an unrealistically short half-life of proteins (on the order of 1ms)^34^. Recently, a biologically feasible filtering mechanism, the spatially coordinated bistable phosphoswitch of PER, was suggested^8^. That is, when enough PER molecules are accumulated in the perinucleus, the bistable switch-like phosphorylation of PER is triggered. As a result, PER molecules at the perinucleus are synchronously phosphorylated, which is necessary for their nuclear entry, and enter the nucleus together. This allows thousands of PER molecules that arrive at the perinucleus at different times to enter the nucleus within a narrow time window. However, such filtering effect was investigated only under an ideal condition that does not include critical factors that affect the protein arrival time, such as variation in cytoplasmic congestion level, cell size, and activator amount.

Here, we investigated whether the bistable phosphoswitch of PER can filter the variability in its arrival time distribution under various noise sources: different cytoplasmic congestion levels, cell sizes, and activator amounts. Specifically, we developed a stochastic model simulating the TTFL of PER, where the PER travel time to the perinucleus is described by a time delay distribution. Thus, changing the delay parameters allows us to effectively describe the change in the travel time due to the variation in cytoplasmic congestion and cell size. Using the framework, we found that bistable phosphorylation of PER can effectively filter the heterogeneity in PER arrival time, resulting in precise repression timing and robust circadian rhythms. On the other hand, such filtering does not occur when PER phosphorylation occurs via conventional phosphorylation mechanisms, such as linear and ultrasensitive phosphorylations. Furthermore, by integrating our model with a previously measured BMAL1 time series in a single cell^14^, we showed that bistable phosphorylation can lead to the precise timing of nuclear entry and robust circadian rhythms even under the large intra- and inter-daily variation in the amount of the activator proteins. Taken together, the bistable phosphoswitch for nuclear entry can filter heterogeneity in protein arrival time and lead to robust rhythms under diverse environments. Our approach sets the stage for systematically exploring how the circadian clock can filter diverse spatiotemporally generated noises in the cell to generate robust circadian rhythms.

## Results

### Synchronous nuclear entry of PER molecules compensates for their spatiotemporal heterogeneity and leads to robust circadian rhythms

In the TTFL of the mammalian circadian clock, the translated PER protein at the cytoplasm (*R*_*c*_) approaches the perinucleus while it forms a complex with CRY proteins and CK1*δ*/*ϵ* (*R*_*p*_), and it is phosphorylated at multiple sites (Figure 1A)^35-39^. Then, the phosphorylated PER complex in the perinucleus 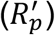 can enter the nucleus, where the PER complex (*R*_*n*_) represses its own transcriptional activator (*A*). The time it takes for PER protein to reach the perinucleus (*τ*) is affected by various biological factors such as cell size (Figure 1B), congestion level (Figure 1C), and the mode of PER transport toward the nucleus (Figure 1D).

**Figure 1.**
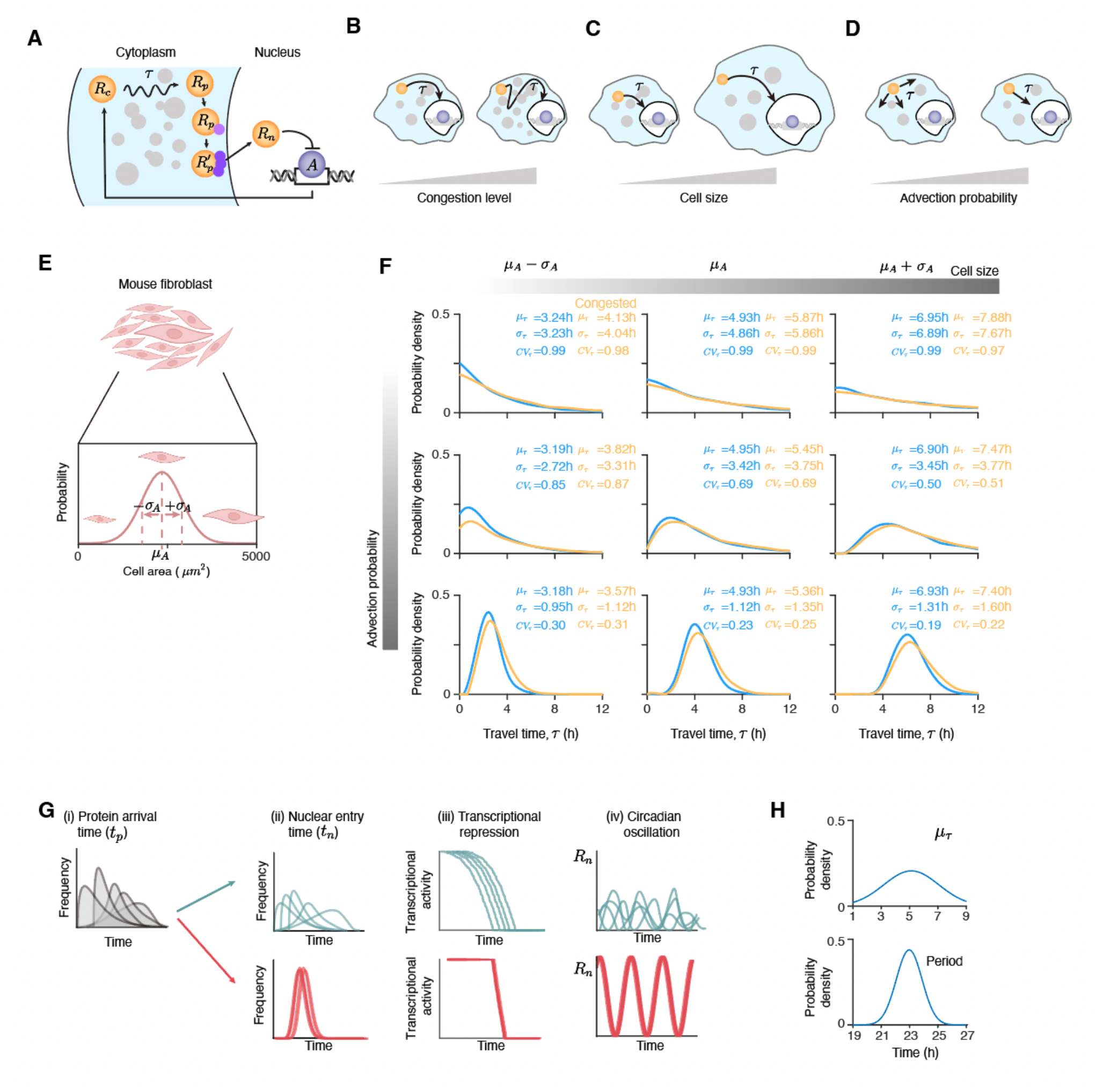
Precise timing of repression is required to generate robust circadian rhythms in various environments. **(A)** Schematic diagram of the circadian clock model. PER protein is synthesized in the cytoplasm (*R*_*c*_), and then it transits toward the perinucleus while it forms a complex with CRY and CK1*δ*/*ε* (*R*_*p*_), and it undergoes multisite phosphorylation. After being phosphorylated, PER complex in the perinucleus 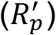 can enter the nucleus (*R*_*n*_) to inhibit its own transcriptional activator (*A*). *R* passes through a cytoplasm, and then arrives at the perinucleus after the travel time *τ*. **(B-D)** The travel time *τ* changes as the crowdedness of cells (B), the size of cells (C) and the chance of advection toward the nucleus (D) change. **(E)** Mouse fibroblast cell size is highly variable, having a mean ± SD, 2346 ± 558 *μm*^2^ (CV=24%)^40^. Thus, the travel time (*τ*) is expected to be highly variable from cell to cell. **(F)** The distribution of travel time (*τ*) varies greatly depending on cytoplasmic congestion level, cell size, and advection probability. As the cell size increases, *μ*_*τ*_ and *σ*_*τ*_ increase. Similarly, a congested cell (yellow) has higher *μ*_*τ*_ and *σ*_*τ*_ than a less congested cell (blue). Increasing advection probability while retaining the same *μ*_*τ*_ leads to a decrease in *CV*_*τ*_. **(G)** Due to the various travel times (*τ*) and the synthesis of proteins over several hours, the arrival time of proteins at the perinucleus (*t*_*p*_) is highly variable. Without a filtering mechanism, the distribution of nuclear entry time (*t*_*n*_) also becomes wide (green line, (ii)), leading to a gradual decline of transcriptional activity (green line, (iii)), and thus weak circadian rhythms (green line, (iv)). A filtering mechanism narrowing the distribution of *t*_*n*_ (red line, (ii)) is needed for sharp transcriptional repression (red line, (iii)), and thus robust circadian rhythms (red line, (iv)). **(H)** Indeed, for mouse fibroblast cells with different sizes^40^, compared to the distribution of mean travel time, the distribution of circadian periods is much narrower, indicating the presence of a noise filtering mechanism.

To quantitatively study how *τ* varies due to these factors, we used an agent-based model that simulates the movement of individual PER molecules from the cell membrane to the perinucleus (see Supplementary Information for details). The movement of PER can be either purely diffused, with a random direction, or directed towards the nucleus in the model. When PER is purely diffused, we used the experimentally measured diffusion coefficient^11,12^. As the advection probability increases, the travel time of PER toward the perinucleus (*τ*) becomes shorter (Figure S1). Thus, as the advection probability increases, we decreased the diffusion coefficient so that the mean of *τ* (*μ*_*τ*_) is maintained. For the simulation, we used the cell size obtained from the average of a mouse fibroblast cell population distribution (*μ*_*A*_)^40^ (Figure 1E) (see Supplementary Information for details). We also increased or decreased the cell size by the standard deviation of the distribution (*σ*_*A*_) to investigate the effect of cell size on *τ*. Furthermore, to consider the effect of congestion level, we also introduced obstacles in the cytoplasm, slowing down the movement of PER, with varying numbers (i.e., different levels of crowdedness).

As cell size increases (left to right, Figure 1F) or crowdedness increases (blue to yellow, Figure 1F), the travel time to reach the perinucleus (*τ*) becomes longer and less precise, i.e., both the mean (*μ*_*τ*_) and standard deviation (*σ*_*τ*_) of the time delay increase. As the chance of advection increases (top to bottom Figure 1F), *CV*_*τ*_ decreases, indicating a more precise arrival time with advection compared to diffusion. However, even with strong advection, the distribution of *τ* remains quite wide and becomes wider as a cell grows and becomes more crowded. Note that the distribution is obtained under the assumption that PER proteins are translated at the same time and the same location. Thus, the distribution of PER arrival times at the perinucleus (*t*_*p*_) is expected to be even more heterogeneous because PER proteins are translated at different places in the cytoplasm over several hours, leading to different travel distances and departure times, respectively^8,11,12^. If 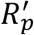 enters the nucleus in the order of arrival at the perinucleus, nuclear entry occurs within a wide time window (i.e., the distribution of *t*_*n*_ is wide) (Figure 1G (ii), green), and thus transcriptional activity decreases gradually (Figure 1G (iii), green), weakening the circadian rhythms (Figure 1G (iv), green)^41-43^. Therefore, to generate strong circadian rhythms, a mechanism narrowing the distribution of *t*_*n*_ is required (Figure 1G, red). The presence of such a filtering mechanism is supported by the distribution of circadian periods across a mouse fibroblast cell population. Specifically, since these cells have highly variable sizes (mean ± SD, 2346 ± 558 *μm*^2^ ; CV=24%) (Figure 1E)^40^, individual cells are expected to have varying protein travel time distributions. Indeed, the calculated distribution of *μ*_*τ*_ across the fibroblast population with a partial differential equation equivalent to the ABM is highly variable (mean ± SD of 5.12 ± 1.94 h) (Figure 1H) (see Supplementary Information for details). However, compared to the variation in travel times, the variation in the circadian periods of the mouse fibroblast cells with different sizes was much smaller (mean ± SD of 22.97 ± 0.91h) (Figure 1H). This indicates the presence of a mechanism to filter the heterogeneous cytoplasmic trafficking time (*τ*) to generate similar circadian periods across different cell sizes. Such a mechanism is expected to be based on the phosphorylation mechanism of *R*_*p*_ because *R*_*p*_ can enter the nucleus only after phosphorylation^3,35,44^.

### Bistable phosphorylation of PER is required to enter the nucleus in the circadian clock

PER protein has multiple CK1-dependent phosphorylation sites where the phosphorylation of these sites occurs in a cooperative manner^8,45^. As a result, the phosphorylation occurs collectively, i.e., the phosphorylation is affected by the local concentration of the PER complex (Figure 2A, red)^8^. Specifically, when the local concentration of the PER complex is lower than the switch-on threshold (Figure 2A, black triangle), phosphorylation does not occur. However, once the local concentration reaches the switch-on threshold, the majority of the PER complex is synchronously phosphorylated, resulting in a sharp increase in the phosphorylated fraction. Furthermore, the high fraction of phosphorylated PER complex persists for awhile, even after the local concentration decreases below the switch-on threshold, until it reaches the switch-off threshold (Figure 2A, gray triangle). To investigate the advantage of having such bistable phosphorylation of *R*_*p*_ in reducing the heterogeneity in the distribution of *t*_*n*_, we compare it with the other typical phosphorylation mechanisms: ultrasensitive phosphorylation and linear phosphorylation. Ultrasensitive phosphorylation also has a switch-on threshold for synchronous phosphorylation, similar to bistable phosphorylation, but it does not have a distinct switch-off threshold (Figure 2A, blue)^43,46^. Thus, the fraction of the 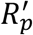 monotonically increases as the total amount of PER in the perinucleus increases, with the sigmoidal pattern having a steep increase near the switch-on threshold. In linear phosphorylation, the fraction of the 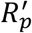 is constant (Figure 2A, green): only the fixed portion of the PER complex is phosphorylated regardless of the local concentration of PER complex.

**Figure 2.**
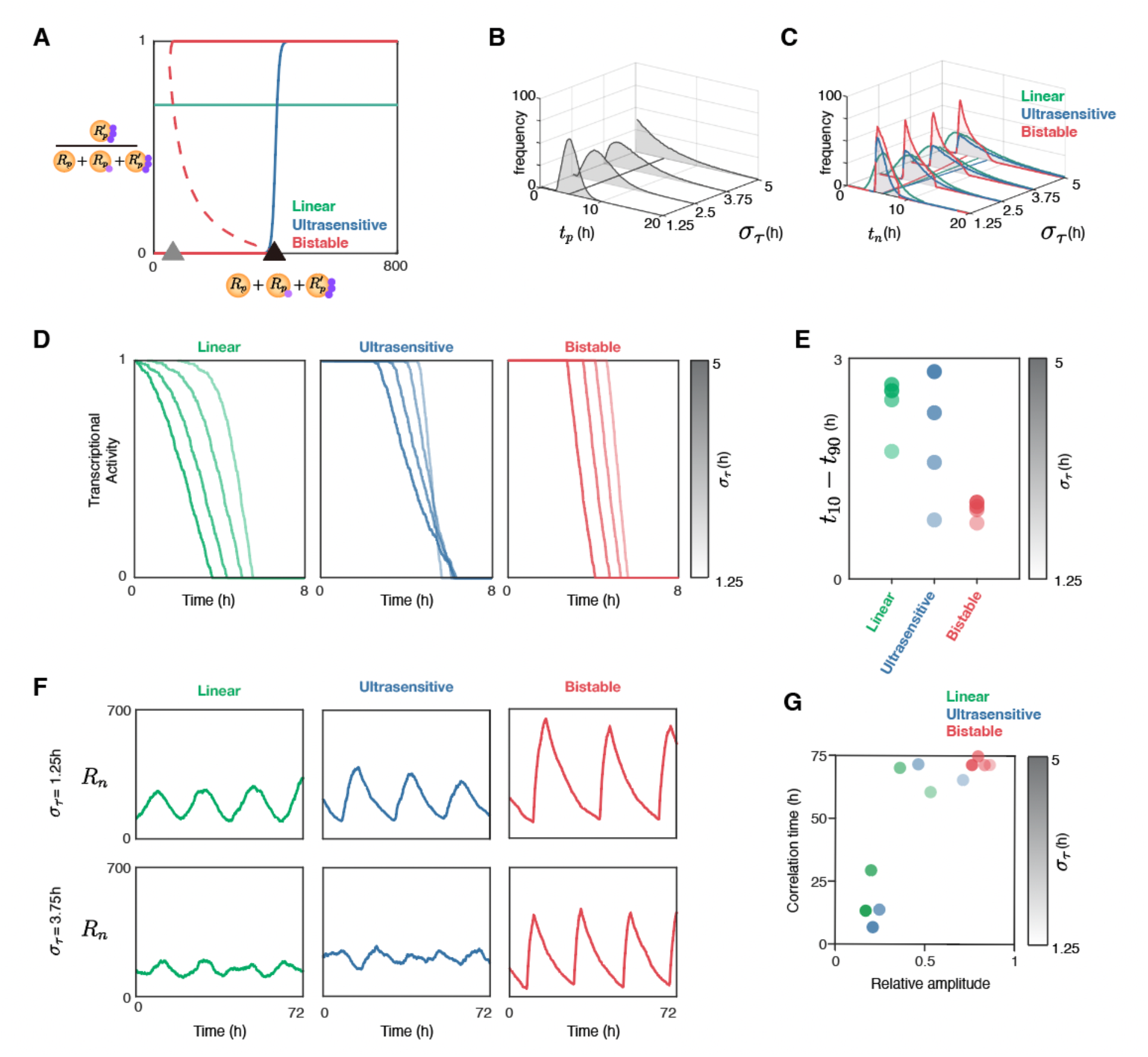
Spatially coordinated collective phosphorylation of PER proteins leads to robust circadian oscillations under noisy protein arrival. **(A)** Three representative phosphorylation mechanisms for nuclear entry. In linear phosphorylation, the fraction of 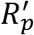 is the same regardless of the local concentration of the total PER complex in the perinucleus. In ultrasensitive and bistable phosphorylations, the fraction of 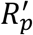 steeply increases when the local concentration of the total PER complex reaches the switch-on threshold (black triangle). With bistable phosphorylation, the fraction of 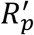 persists even when the local concentration of the total PER complex decreases below the switch-on threshold until the local concentration of the total PER complex reaches the switch-off threshold (gray triangle). **(B-C)** As the protein travel time becomes noisy (i.e., *σ*_*τ*_ increases), thousands of PER molecules arrive at the perinucleus within a wide time window (*t*_*p*_). As a result, the distribution of nuclear entry time (*t*_*n*_) also becomes significantly wider with linear (green) or ultrasensitive (blue) phosphorylation, but not with bistable phosphorylation (red). Here, the distributions were obtained from 200 repeated simulations. **(D)** As *σ*_*τ*_ increases (light to dark colors), with linear and ultrasensitive phosphorylations, transcriptional activity decreases gradually (green and blue). In contrast, with bistable phosphorylation, transcriptional activity decreases sharply regardless of *σ*_*τ*_ (red). **(E)** The sensitivity of transcription repression is quantified by measuring the time taken for the transcriptional activity to decrease from 90% (*t*_90_) to 10% of the maximal transcriptional activity (*t*_10_). The smaller value of *t*_10_ − *t*_90_ indicates the sharper repression of transcription. **(F)** Robust oscillations are maintained with bistable phosphorylation even under noisy cytoplasmic trafficking (i.e., high *σ*_*τ*_), but not with linear and ultrasensitive phosphorylations. **(G)** The robustness of the oscillations was quantified by the correlation time and the relative amplitude of *R*_*n*_ oscillatory time series. The longer correlation time indicates the more robust oscillation.

### Bistable phosphorylation of PER allows sharp transcriptional repression despite the heterogeneous PER arrival times at the perinucleus

We compared how the distribution of nuclear entry times (*t*_*n*_) is affected as the variance in the arrival time (*t*_*p*_) increases depending on the three phosphorylation mechanisms regulating nuclear entry. For this, we constructed a mathematical model which describes part of the TTFL (Figure 1A): the cytoplasmic trafficking of *R*_*c*_ to *R*_*p*_, the phosphorylation of *R*_*p*_ to 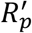, and the nuclear entry of 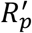 to *R* (see STAR Methods for detailed model descriptions and Table S1 for the propensity functions of reactions). To focus on how phosphorylation affects the distribution of *t*_*n*_, we assumed that PER molecules are not degraded. Additionally, the time spent during the cytoplasmic trafficking was assumed to be gamma distributed, because the distributions of *τ* in Figure 1F approximately follow the gamma distribution. After arrival at the perinucleus, *R*_*p*_ undergoes phosphorylation with one of the three phosphorylation mechanisms (Figure 2A) and enters the nucleus. We assumed that *R*_*p*_ stays at the perinucleus and that the phosphorylation of the PER protein only occurs at the perinucleus, based on a previous study^8^, which observed the enrichment of the PER protein and phosphorylation mainly at the perinucleus. To describe the different types of phosphorylation, we adopted the modular approach, which describes complex multistep reactions with a single phenomenological module that qualitatively describes the dynamical behavior^46,47^. This allowed us to simulate the complex phosphorylation process, which is necessary for nuclear entry (see STAR Methods for details). We simulated the situation where 1,000 PER complexes travel from the cell membrane to the perinucleus using the delayed Gillespie algorithm^48,49^. This allowed us to calculate the time when each *R*_*c*_ arrives at the perinucleus (*t*_*p*_) and then when it enters the nucleus (*t*_*n*_). In this way, we obtained the distribution of *t*_*p*_ and *t*_*n*_ of the 1,000 PER complexes, which quantifies the heterogeneity in the protein arrival time and the nuclear entry time, respectively.

As we increase the standard deviation of the travel time (*σ*_*τ*_) while keeping its mean (*μ*_*τ*_) constant, the distribution of *t*_*p*_ becomes wider (Figure 2B). As a result, the distribution of *t*_*n*_ also becomes wider with linear and ultrasensitive phosphorylations (Figure 2C, green and blue), and thus transcriptional activity decreases gradually (Figure 2D, green and blue). However, the narrow distribution of *t*_*n*_ is maintained with bistable phosphorylation (Figure 2C, red), and thus transcriptional repression occurs sharply regardless of the *σ*_*τ*_ (Figure 2D, red). To quantify the sensitivity of the transcriptional repression (i.e., how rapidly the transcriptional activity is decreased), the difference between the time when the transcriptional activity reaches 10% of the maximal transcriptional activity (*t*_10_) and the time when the transcriptional activity reaches 90% of the maximal transcriptional activity (*t*_90_) was calculated. With bistable phosphorylation, *t*_10_ − *t*_90_ is small (i.e., sharp repression) regardless of *σ*_*τ*_ (Figure 2E). On the other hand, with linear and ultrasensitive phosphorylations, *t*_10_ − *t*_90_ greatly increases as *σ*_*τ*_ increases. This indicates that bistable phosphorylation, but not linear and ultrasensitive phosphorylations, allows sharp transcriptional repression even when the protein arrives at the perinucleus with heterogeneous timing (i.e., the variance of *t*_*p*_ is large.)

### Robust circadian rhythms can be generated even with highly variable travel time through bistable phosphorylation

In the previous section, to focus on the nuclear entry time (*t*_*n*_) of 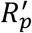, we used a model describing part of the TTFL (Figure 1A). Now, we have extended the model to fully describe the TTFL. Specifically, in the model, the transcriptional activator (*A*) is suppressed by *R*_*n*_ via protein sequestration^19,21,22,28^, which closes the negative feedback loop. Furthermore, the degradations of *R*_*c*_, *R*_*p*_, 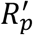, and *R*_*n*_ were also incorporated into the model. With this model, we investigated how the noise level in the travel time (i.e., *σ*_*τ*_) affects circadian rhythms. As *σ*_*τ*_ increases, the simulated oscillations of *R*_*n*_ become noisy with linear and ultrasensitive phosphorylations (Figure 2F, green and blue). On the other hand, even when *σ*_*τ*_ increases, robust oscillations are maintained with bistable phosphorylation (Figure 2F, red). To quantify the accuracy and the strength of the simulated oscillations, we calculated the correlation time and the relative amplitude of the time series of *R*_*n*_ (see STAR Methods). The correlation time describes how fast the autocorrelation function of the oscillatory time series decays^50,51^. If the time series exhibits a noisy oscillation, autocorrelation will decay rapidly, resulting in a short correlation time. The correlation time becomes short and relative amplitude becomes small with linear and ultrasensitive phosphorylations when *σ*_*τ*_ increases (Figure 2G, green and blue). In other words, linear and ultrasensitive phosphorylations become less likely to generate rhythms as *σ*_*τ*_ increases. This result is consistent with a previous study^52^, which showed that negative-feedback oscillators find it challenging to generate rhythms with a distributed delay that has a large width. On the other hand, the correlation time is long and the relative amplitude is large regardless of *σ*_*τ*_ with bistable phosphorylation (Figure 2G, red). Taken together, with bistable phosphorylation, robust oscillations can be generated even when *σ*_*τ*_ is large.

### The effect of travel time on circadian periods is reduced by bistable phosphorylation

Next, we examined if bistable phosphorylation can generate robust and precise rhythms even when *μ*_*τ*_ and *σ*_*τ*_ change simultaneously. For this, we simulated the TTFL model used in Figures 2F and 2G for cells with different *μ*_*τ*_ and *σ*_*τ*_. We used *μ*_*τ*_ obtained from 1,000 mouse fibroblast cells with different sizes (Figure 1H), and fixed *CV*_*τ*_ to be 0.5 so that *σ*_*τ*_ = 0.5 · *μ*_*τ*_. As *μ*_*τ*_ and *σ*_*τ*_ vary, bistable phosphorylation leads to a much narrower distribution of periods (SD=1.97h) compared to linear and ultrasensitive phosphorylations (SD=4.37h and SD=4.41h, respectively) (Figure 3A). Even when we fixed *CV*_*τ*_ to different values (such as 0.25 or 0.75), we could obtain consistent results (Figure S2). Thus, bistable phosphorylation generates robust and accurate rhythms when both the mean (*μ*_*τ*_) and the standard deviation (*σ*_*τ*_) vary.

**Figure 3.**
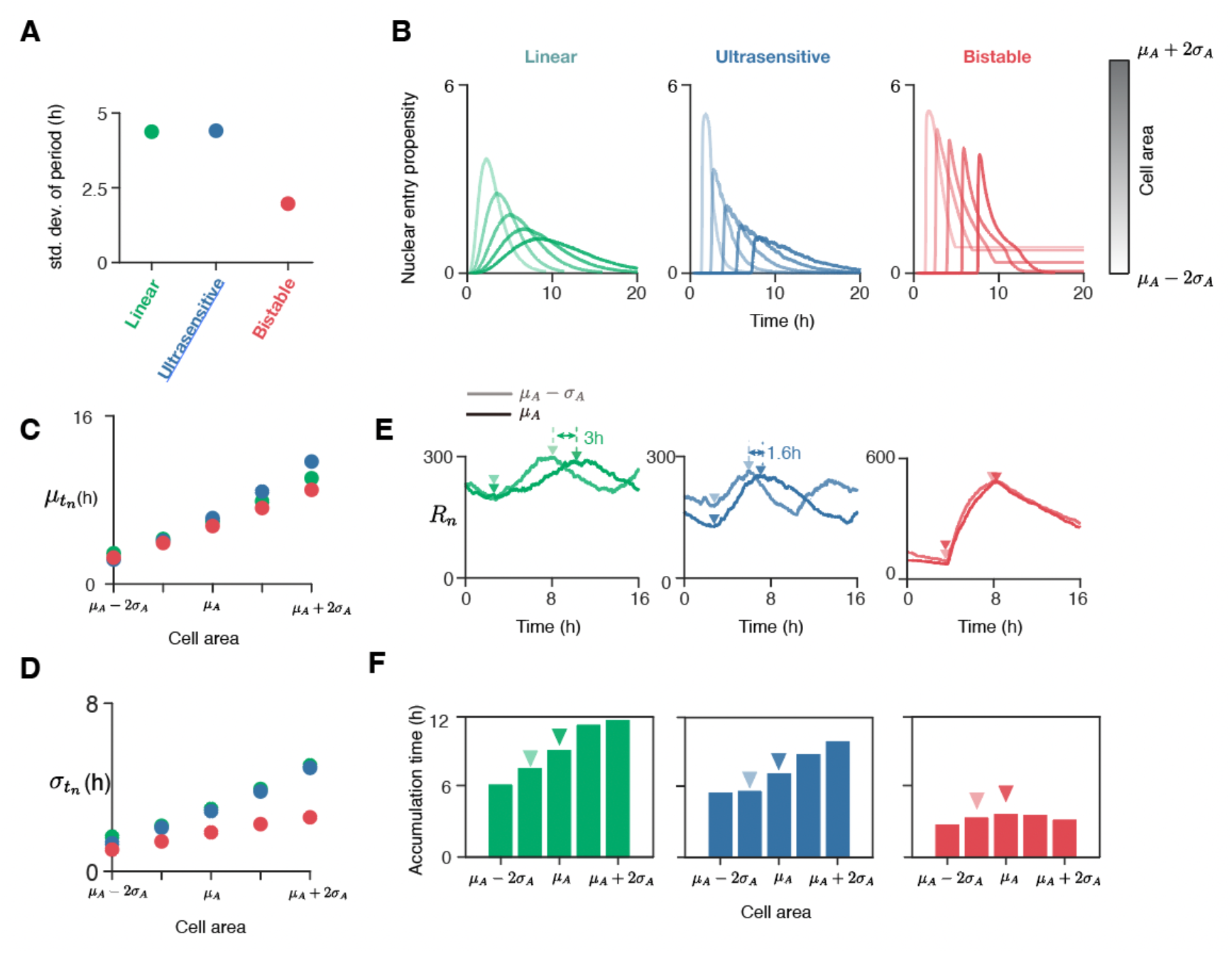
Bistable phosphorylation enables cells of different sizes to have similar periods. **(A)** For mouse fibroblast cells with different sizes^40^, although the standard deviation of the mean (*μ*_*τ*_) and the standard deviation (*σ*_*τ*_) of time spent during the cytoplasmic trafficking are ∼2h, the standard deviation of the circadian periods is only a half of them (SD=0.91h). **(B)** The standard deviation of the simulated circadian periods for 1,000 cells, whose areas were sampled from the *N*(2346*μm*^2^, 558^2^*μm*^4^). **(C)** In a larger cell, *μ*_*τ*_ and *σ*_*τ*_ increase. Thus, the time trajectories of the nuclear entry propensity (Tables S1 and S2) are shifted to the right and become wider. Note that with bistable phosphorylation, the nuclear entry occurs in a narrow time window, even in larger cells. The distributions were obtained with 200 repeated simulations. *μ*_*A*_ (=2346*μm*^2^) and *σ*_*A*_ (=558*μm*^2^) are the mean and standard deviation of fibroblast cell area, respectively. **(D-E)** While the mean of the nuclear entry time 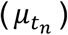 increases similarly in all three phosphorylation mechanisms, bistable phosphorylation resulted in a far smaller standard deviation in nuclear entry time 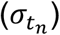 than linear and ultrasensitive phosphorylations. **(F-G)** As a result, the length of time during which *R*_*n*_ accumulates from the trough (triangle) to the peak (triangle) of *R*_*n*_ does not change regardless of cell area with bistable phosphorylation, but this is not the case with linear and ultrasensitive phosphorylations.

We then investigated how bistable phosphorylation compensates for the effect of change in *μ*_*τ*_ and *σ*_*τ*_ on the circadian period. To do this, a model describing part of the TTFL (Figures 2B and 2C) with different *μ*_*τ*_ and *σ*_*τ*_ was utilzed. We used five representatives of *μ*_*τ*_ obtained from five cell areas: *μ*_*A*_ − 2*σ*_*A*_, *μ*_*A*_ − *σ*_*A*_, *μ*_*A*_, *μ*_*A*_ + *σ*_*A*_, *and μ*_*A*_ + 2*σ*_*A*_ (Figures 1E and 1H top). We again set *σ*_*τ*_ = 0.5 · *μ*_*τ*_ by fixing 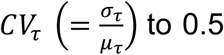 as in Figure 3A. We then examined how these various values of *μ*_*τ*_ and *σ*_*τ*_, obtained from five different cell areas, affected the nuclear entry time (*t*_*n*_) under each phosphorylation mechanism. As the cell size increases, and thus *μ*_*τ*_ increases, the simulated time trajectories of the nuclear entry propensity (Tables S1 and S2) are shifted to the right (Figure 3B). Thus, the mean of nucleus entry time (*t*_*n*_) increases for all three phosphorylation mechanisms (Figure 3C), which is expected to lengthen the period. Furthermore, as *σ*_*τ*_ also increases due to the cell size increase, PER molecules enter the nucleus in a wider time window, and thus the time trajectories of the nuclear entry propensity become wider and lower. Lower nuclear entry propensity results in a slow increase in *R*_*n*_ (Figure 3B), which may further lengthen the period. Interestingly, such change is much smaller with bistable phosphorylation compared to linear and ultrasensitive phosphorylation (Figure 3B). That is, the increase in the standard deviation of *t*_*n*_ is significantly larger with linear and ultrasensitive phosphorylations than with bistable phosphorylation (Figure 3D). As a result, when the TTFL model was simulated for each phosphorylation mechanism, the time between the minimum of *R*_*n*_ and the maximum of *R*_*n*_ became longer (i.e., *R*_*n*_ accumulated more slowly) in a larger cell with linear and ultrasensitive phosphorylations, but not with bistable phosphorylation (Figure 3E). Specifically, the mean length of time during which *R*_*n*_ increased more than two-fold change varied from ∼4h to ∼10h as the cell area increased from *μ*_*A*_ − 2*σ*_*A*_ to *μ*_*A*_ + 2*σ*_*A*_ with linear and ultrasensitive phosphorylations (Figure 3F, green and blue). On the other hand, with bistable phosphorylation, the accumulation time of *R*_*n*_ was nearly the same regardless of the cell area (Figure 3F, red). This explains why the change in the circadian period due to the cell size variation is smaller with bistable phosphorylation than with other phosphorylation mechanisms (Figure 1H).

### Bistable phosphorylation enables robust circadian rhythms despite noisy activator rhythms

In the previous sections, the amount of activator protein was assumed to be constant over time. However, the amount of activator molecules, such as BMAL1, oscillates with a period of ∼24h^13,14,35,53^ (Figure 4A). In addition, the amplitude and the peak-to-peak period of BMAL1 protein fluctuate from day to day. Specifically, the amplitude and the peak-to-peak period of experimentally measured BMAL1 proteins in a single cell highly vary^14^: 286±35 and 25.4±5.7h (Figure 4A). Furthermore, an additional ‘bump’ often exists between the peaks. Due to the noisy pattern, it is expected that the synthesis timing of PER will be heterogeneous, resulting in the heterogeneous protein arrival time distribution. We investigated whether bistable phosphorylation enables robust circadian rhythms of PER even under such fluctuation in activators, resulting in noisy transcription of PER. For this, we simulated the TTFL model (Figure 1A), where the activator level changes according to the experimentally measured BMAL1 (Figure 4A). The amplitude of oscillation is highly variable with linear and ultrasensitive phosphorylations (Figure 4B, green and blue). Furthermore, additional peaks in the nuclear PER complex (*R*_*n*_) are observed with linear and ultrasensitive phosphorylations. In contrast, with bistable phosphorylation, robust rhythms with similar amplitudes are generated (Figure 4B, red). Moreover, the additional peaks do not appear. As a result, the CV of the peak-to-peak period and the amplitude are much smaller with bistable phosphorylation than with linear and ultrasensitive phosphorylations (Figure 4C). Taken together, bistable phosphorylation enables robust circadian rhythms despite fluctuation in activator rhythms.

**Figure 4.**
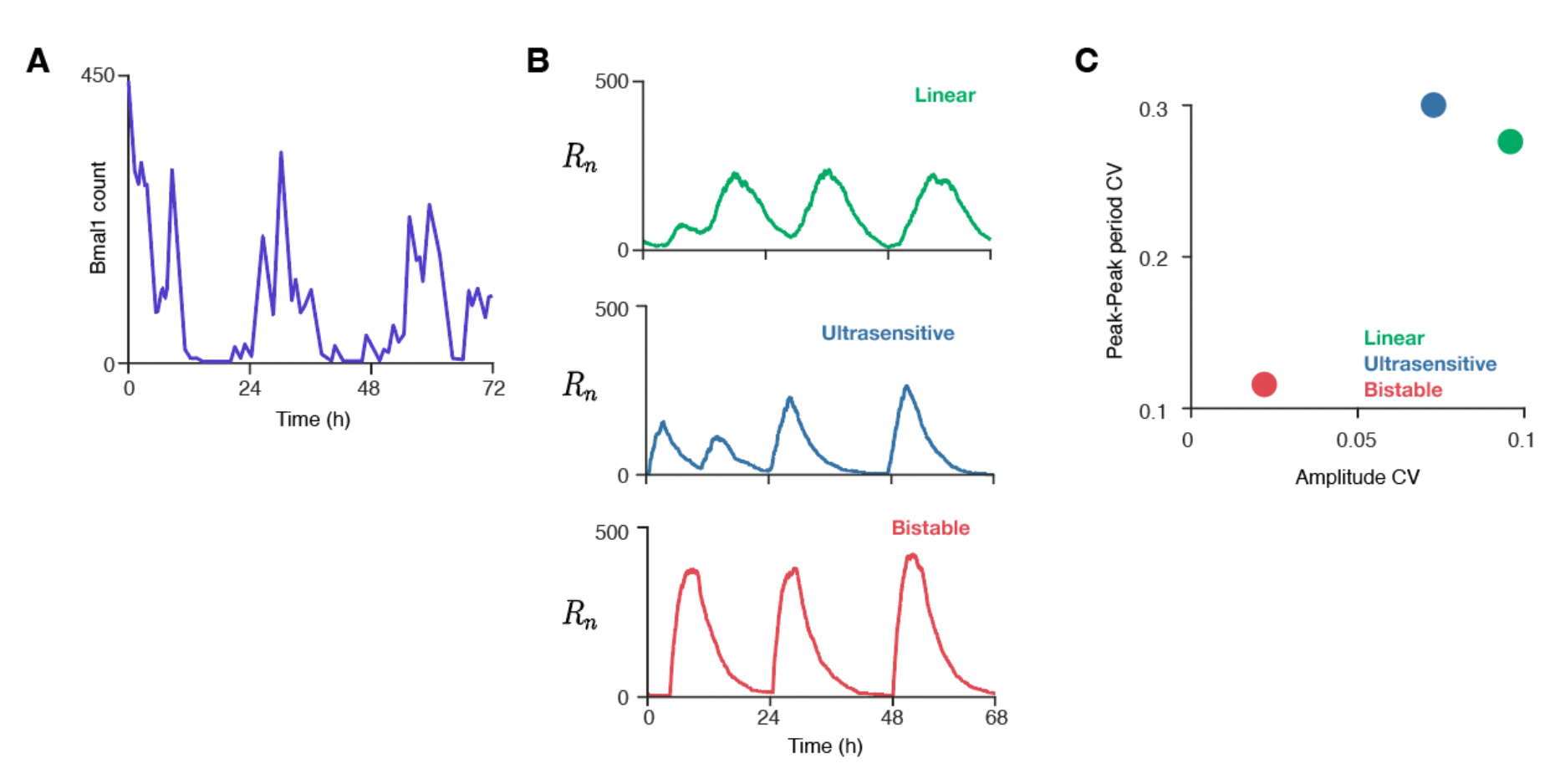
Bistable phosphorylation enables cells to sustain robust circadian rhythms even under noisy activator rhythms. **(A)** The level of activator protein BMAL1 fluctuates across a day and exhibits daily change. In addition, an additional bump often exists, resulting in more than one peak during a single cycle. The graph was retrieved from Blanchoud et al.^14^. **(B)** Even with such noisy activator time series, bistable phosphorylation leads to robust rhythms, unlike linear and ultrasensitive phosphorylations. **(C)** As a result, the CV of both the peak-to-peak period and the amplitudes is small with bistable phosphorylation, but not with linear and ultrasensitive phosphorylation.

## Discussion

For the TTFL to generate robust circadian rhythms, precise nucleus entry of the PER complex is critical^7-10^. However, the distribution of the PER complex arrival time at the perinuclear region is highly heterogeneous due to various noise sources. Despite this, precise nucleus entry of the PER complex and transcriptional repression occur in the circadian clock. To investigate the molecular mechanism underlying the unexpected precision of the circadian clock, we constructed a mathematical model describing the effect of protein arrival time on circadian rhythms (Figure 1A). We found that when the protein arrival time varies due to the change of cytoplasmic congestion level, cell size, and activator level, bistable phosphorylation, but not linear and ultrasensitive phosphorylations, leads to precise repression timing, and thus robust circadian rhythms (Figures 2-4). This indicates that the bistable phosphoswitch is the key mechanism to filter the spatiotemporal noise in the cell to generate circadian rhythms. Spatiotemporally regulated bistable phosphorylation can also play a critical role in other biological oscillators, such as the cell cycle^54-56^.

The mathematical model used in our study describes the spatiotemporal dynamics of the TTFL of the circadian clock. In previous studies, the spatiotemporal dynamics of molecules from the cytoplasm to the nucleus were directly described using PDE^33^ or agent-based models^8,32^. This approach was computationally expensive and not flexible to describe the various noise sources for the protein arrival time of molecules to the perinucleus. To describe the spatiotemporal dynamics of molecules simply and flexibly, we used a travel time distribution as in previous studies, which described a chain of signaling processes or protein synthesis^57-61^. Furthermore, it is challenging to model ultrasensitive and bistable phosphorylations since they are generated from complex combinations of multiple reactions. To resolve this, we utilized a modular approach that describes those complex multistep reactions with a phenomenological module, which was developed in work by De Boeck et al.^46^. This allowed us to effectively capture the key dynamics of the phosphorylation mechanisms without comprehensively modeling the underlying complex reactions. Using the distributed travel time and the modular approach are effective tools for describing complex intracellular spatiotemporal dynamics.

The nuclear entry of PER is delayed by the bistable phosphoswitch until a certain accumulation level is reached. Such delayed nuclear entry of PER has been observed in previous studies with *Drosophila* and mammals^8,62^. However, a study reported a rapid PER import when a protein synthesis inhibitor, cycloheximide (CHX), was applied and then removed from the cell^63^. The cause for the conflicting results observed in previous studies remains unclear, as the detailed mechanisms for nuclear entry and its rate constants have not been fully explored. One potential explanation is that CHX disrupts the phosphoswitch by inhibiting the synthesis of proteins, including kinase and phosphatase. Note that the balance between kinase and phosphatase is essential for the bistable phosphoswitch function^8,64^. To investigate the nuclear entry of PER regulated by the bistable phophoswitch, it is important to avoid perturbing the endogenous condition. This is also important for investigating the nuclear entry in biological systems other than the circadian clock. For example, in pancreatic *β*-cells, the nuclear entry of cPKA should be slow to distinguish the time scales of external signals^65^. While the underlying molecular mechanisms of such slow nuclear entry have not been known, the bistable phosphoswitch could be a potential molecular mechanism leading to such regulation of cPKA in pancreatic *β*-cells.

Alzheimer’s disease (AD) has been reported to disrupt the circadian clock^66^. Such distribution appears to be due to tauopathy, which triggers the formation of neurofibrillary tangles, and thus increases cytoplasmic crowdedness^8^. However, even though tauopathy is one of the earliest events in AD development^67^, the activity rhythms of mild symptomatic dementia patients are not significantly different from those of non-patients^68,69^. This can be explained by our results: the bistable phosphoswitch allows the circadian rhythms to function normally up to a reasonable increase in intracellular congestion level.

### Limitations of the study

In this work, we utilized an extension of the Gillespie algorithm to simulate the system describing the TTFL. This allowed us to describe the effects of various noise sources on circadian rhythms. To use the Gillespie algorithm for a system containing non-elementary propensity functions (e.g., Hill functions) other than functions from mass-action kinetics, timescale separation is necessary^70-72^. Thus, we assumed that the synthesis and the degradation of PER are slower than other reactions, such as binding and unbinding of PER to activator proteins and each step of its phosphorylation and dephosphorylation. However, timescale separation is often not enough for accurate stochastic simulations, unlike deterministic simulations; i.e., the condition for using the non-elementary propensity functions for stochastic simulation is stricter than the deterministic simulations^71,73-75^. Furthermore, the validity of using non-elementary propensity functions for stochastic simulations in the presence of distributed delays has not been investigated, which will be interesting in future work.

Clock proteins have complex interactions with each other. For example, BMAL1 regulates PER2 transcription, while PER2 regulates BMAL1 transcription by affecting the transcription of REV-ERB and RORs, which are transcriptional regulators of BMAL1^4^. However, our model omits the regulation of BMAL1 by PER2 for simplicity in Figure 4. Further work could include regulation from PER2 to BMAL1 and examining the interaction between the repressor proteins and activator proteins in the circadian clock.

Cell size keeps changing due to cell growth or cell division, which affects the TTFL of the circadian clock. Specifically, as cell size changes, the cytoplasmic trafficking of PER molecules changes, which is investigated in our study by using a time delay distribution (Figures 1F, 3A-3F). However, we did not investigate other critical factors, such as the dilution of the PER molecules during cell growth and their partition into daughter cells after cell division^76-78^. It would be interesting in future work to extend our study to incorporate dilution due to cell growth and the partition of molecules after cell division to investigate further the effect of the cell size changes on circadian rhythms.

## Acknowledgements

This work was supported by the Human Frontiers Science Program Organization (grant no. RGY0063/2017) (J.K.K.), the Institute for Basic Science (grant no. IBS-R029-C3) (J.K.K.), the Korea University Grant (S.L.) and the National Research Foundation of Korea (NRF) grant funded by the Korea government (MSIP) (No. 2020R1A2C1A01100114) (S.L).

## Author contributions

All authors designed the study. S.J.C. performed and D.W.K., S.L. and J.K.K. contributed to computational modeling and simulation. All authors analyzed the data. J.K.K. supervised the project. S.J.C. and J.K.K. wrote the draft of the manuscript, and all authors revised the manuscript.

## STAR★Methods

### RESOURCE AVAILABILITY

#### Lead contact

Further information and requests for resources and reagents should be directed to and will be fulfilled by the Lead Contact, Jae Kyoung Kim.

#### Materials availability

This study did not generate new unique reagents.

#### Data and code availability

- This paper analyzes publicly available existing data. These accession numbers for the datasets are listed in the key resources table.
- The MATLAB and Netlogo codes of the computational package are available in the following Database: The link will be available upon acceptance of the manuscript.
- Any additional information required to analyze the data is available from the lead contact upon request.

## METHODS DETAILS

### The stochastic model of the circadian clock simulating the TTFL with the spatiotemporal behavior of PER proteins

We extended a previous mathematical model of the mammalian circadian clock^19,21,22,28^ to study the influence of the spatiotemporal behavior of PER proteins on circadian rhythms (see Tables S1 and S2 for detailed reactions and parameters used in the simulation). Specifically, our model consists of four variables: PER protein in the peripheral cytoplasm (*R*_*c*_), unphosphorylated PER complex in the perinucleus (*R*_*p*_), phosphorylated PER complex in the perinucleus 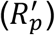, and PER complex in the nucleus (*R*_*n*_), which degrade with the same rate constant of *λ*_*d*_.

In the model, *R*_*c*_ is synthesized at the rate of *λ*_*p*_ max 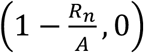, where *A* is the number of activator proteins and *λ*_*p*_ is the maximum synthesis rate. The synthesis rate is proportional to the fraction of free activator that is not sequestrated by PER complex in the nucleus, described by max 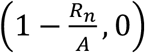. This is an approximated form of the quasi-steady state of the fraction of free activators found by applying the total quasi-steady state approximation reduction to the detailed model describing the circadian clock^19,21,22,28^ under the assumption that PER quickly binds and unbinds to the activator: 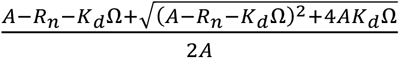, where *K*_*d*_ is the dissociation constant between the activator and PER, and Ω is the volume of the system. Then, by further assuming that the binding is tight (*K*_*d*_ → 0), the fraction of free activator can be approximated by max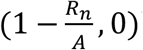. Up to Figure 3, *A* was fixed to be 300, assuming the number of the activator is constant over time to focus on the spatiotemporal dynamics of PER. In Figure 4, to investigate whether bistable phosphorylation enables robust circadian rhythms even under noisy activator rhythms, we changed *A* according to the experimentally measured time series of BMAL1 molecules in a single cell^14^. From 72h-long experimentally measured BMAL1 data, the first four hours were excluded to avoid the effect of the initial shock for the experiment and then repeated to generate a long time series of *A*. Since the experimentally measured BMAL1 time series was discrete, we linearly interpolated the data to obtain the amount of BMAL1 in continuous time.

After being synthesized, *R*_*c*_ passes through the cytoplasm crowded with obstacles during a gamma-distributed travel time *τ*. Specifically, when each *R*_*c*_ molecule is produced, *τ* is sampled from the gamma distribution and assigned to each molecule. Here, the gamma distribution is utilized because it successfully captures the distribution of time spent during the cytoplasmic trafficking. To determine the shape of the gamma distribution, we utilized a previously proposed method using the ABM model, which describes the cytoplasmic trafficking, so that the mean and the standard deviation of *τ* can be obtained (see Supplemental Information for details). When the experimentally measured diffusion coefficient of PER of 0.2*μm*^2^/*s*^11,12^ and the mean area of the fibroblast cell of 2346*μm*^2^ were used, the PDE simulation resulted in *μ*_*τ*_=4.9h. This value of *μ*_*τ*_ was used in Figure 2 and Figure 4. *σ*_*τ*_ was set to be *σ*_*τ*_ = 0.5 · *μ*_*τ*_ to ensure linear and ultrasensitive phosphorylations to generate oscillations.

The PER complex that arrives at the perinucleus (*R*_*p*_) undergoes phosphorylation to be 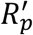. Then, 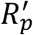 is either dephosphorylated to *R* with the rate of 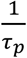 or enters the nucleus with the rate of *λ*_*n*_. To simply describe the phosphorylation based on multiple reactions, we adopted the modular approach^46^, which treated the phosphorylation reaction as a single phenomenological module. That is, the module takes the total amount of PER complex in the perinucleus 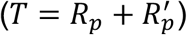 as the input and gives the fraction of the 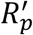 in the perinucleus as the output. Among possible responses that the module can generate, we chose linear phosphorylation, ultrasensitive phosphorylation, and bistable phosphorylation^47^. With linear phosphorylation, the fraction of 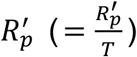 is constant regardless of *T* (Figure 2A, green). Such response is obtained by utilizing the propensity function for the phosphorylation 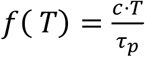, where *c* is between 0 and 1. In our simulation, the time scale of phosphorylation and dephosphorylation reactions, *τ*_*p*_, was set to be sufficiently short 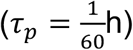 so that the phosphorylation and dephosphorylation quickly equilibrate (i.e.,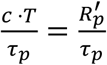). As a result, the fraction of 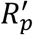 equilibrates to 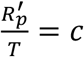. Here, *c* = 0.7 was chosen to make the fraction of 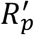 be 70%. For ultrasensitive phosphorylation, the fraction of 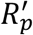 monotonically increases as *T* increases, with a steep increase near the threshold. To describe ultrasensitive phosphorylation, the propensity function 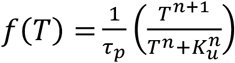 was used. Thus, the fraction of 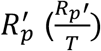 quickly equilibrates to 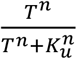, which was used to describe the ultrasensitive response previously^43,46^. We set *K*_*u*_ = 400 and *n* = 70 to obtain a steep increase in the fraction of 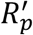 when *T* reaches 400 (Figure 2A, blue). For bistable phosphorylation, the response is bifurcating (i.e., the response qualitatively varies depending on variables other than *T*), being affected by 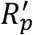. Thus, the response of bistable phosphorylation is described not only by *T*, but also by 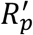. To describe bistable phosphorylation, we used an implicit function 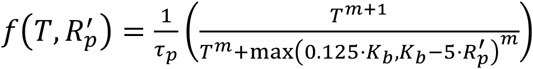 after modifying a function generating an S-shaped response curve (i.e., bistable response)^46^. 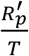 quickly equilibrates to 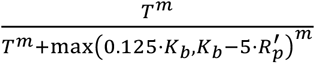, which steeply increases when *T* increases to reach the switch-on threshold, *K*_*b*_. Once *T* reaches *K*_*b*_, the fraction of 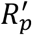 stays near one even though *T* decreases below *K*_*b*_ until it reaches the switch-off threshold, which is about 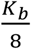. Here, we chose *m* = 30 and *K*_*b*_ = 400 (Figure 2A, red).

The model was simulated using the delayed Gillespie algorithm^48,49^. For the Gillespie simulation, the volume of the system was set to 1. See Tables S1 and S2 for detailed descriptions of propensity functions and parameters used in the simulation, respectively. In Figures 2B-2E and Figures 3B-3D, to focus on how the phosphorylation affects the distribution of *t*_*n*_, the synthesis of *R*_*c*_ and the degradation of PER molecules are excluded from the simulation. Specifically, we set *λ*_*p*_ and *λ*_*d*_ to be 0 to make these reactions not occur. The initial condition was given by *R*_*c*_ = 1000,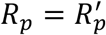 = *R*_*n*_ = 0.

## QUANTIFICATIONS AND STATISTICAL ANALYSIS

### Quantification of features of the oscillation

We calculated several features (e.g., period, amplitude, correlation time) from the oscillatory time series. First, in Figure 2G, we calculated the correlation time^50,51^ and the relative amplitude^31^, as done in previous studies. Specifically, to calculate the correlation time, we estimated parameters *τ*_*C*_ and *P* by fitting 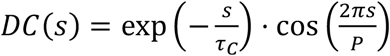 to the autocorrelation function 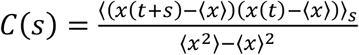 where *x*(*t*) is the *R*_*n*_ time series obtained from the simulation. Then, the correlation time is defined as *τ*_*C*_, and it describes how fast the autocorrelation decays over time. Thus, a more robust oscillation has a larger *τ*_*C*_. The relative amplitude was calculated as the fraction of the amplitude (i.e., the difference between the peak height and trough height of the given rhythms) to the peak height. To calculate the correlation time and the relative amplitude, the *R*_*n*_ time series obtained from the simulation for 100 days (i.e., 2,400h) was used after the first 10 days were excluded to avoid the effect of transient dynamics. We repeated ten simulations with the same initial condition of having no PER molecule in the cell. Then, we took the average correlation time over ten repetitions.

In Figure 3A, we calculated the period of the oscillation by fitting the *C*(*s*) to *DC*(*s*) for simulated time series of *R*_*n*_ for 100 days after excluding first 10 transient days. Then, we used the estimated *P* as the period. We repeated this over ten different simulated trajectories and used the average of the estimated *P* as the period.

To calculate the accumulation time of *R*_*n*_ in Figure 3F and the peak-to-peak period in Figure 4, the time series of *R*_*n*_ simulated for 100 days after excluding first 10 transient days was used. Then, the accumulation time of *R*_*n*_ was measured from each cycle over 10 repeated simulations and then the mean of *R*_*n*_ accumulation time was calculated. The peak-to-peak period was measured from each cycle over 10 repeated simulations, and then CV was calculated (Figure 4).

## KEY RESOURCE TABLE

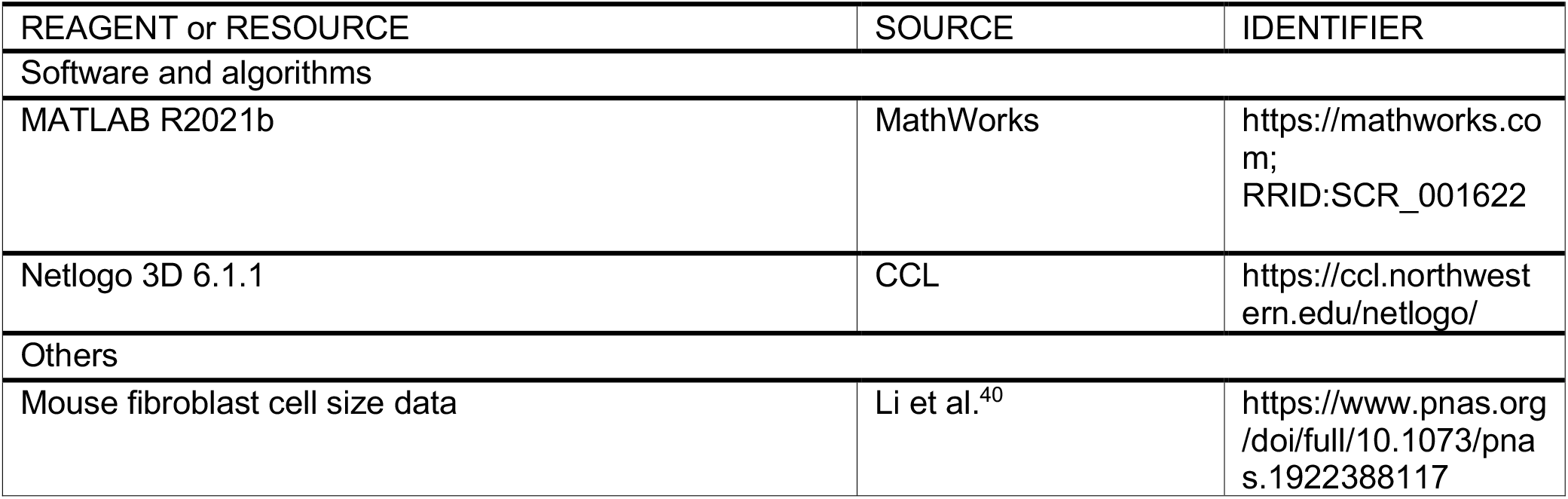
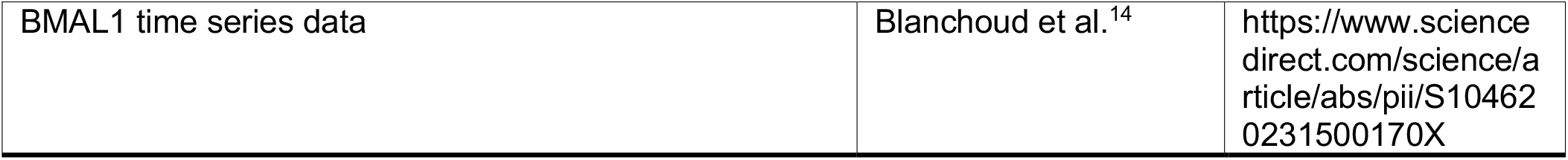

| REAGENT or RESOURCE | SOURCE | IDENTIFIER |
| --- | --- | --- |
| Software and algorithms |  |  |
| MATLAB R2021b | MathWorks | https://mathworks.com;<br>RRID:SCR_001622 |
| Netlogo 3D 6.1.1 | CCL | https://ccl.northwestern.edu/netlogo/ |
| Others |  |  |
| Mouse fibroblast cell size data | Li et al.<sup>40</sup> | https://www.pnas.org/doi/full/10.1073/pnas.1922388117 |
| BMAL1 time series data | Blanchoud et al.<sup>14</sup> | https://www.sciencedirect.com/science/article/abs/pii/S104620231500170X |

